# Scarless removal of large resistance island AbaR results in antibiotic susceptibility and increased natural transformability in *Acinetobacter baumannii*

**DOI:** 10.1101/2020.04.29.067926

**Authors:** Anne-Sophie Godeux, Elin Svedholm, Agnese Lupo, Marisa Haenni, Maria-Halima Laaberki, Xavier Charpentier

**Affiliations:** CIRI, Centre International de Recherche en Infectiologie, Univ Lyon, Inserm, U1111, Université Claude Bernard Lyon 1, CNRS, UMR5308, ENS de Lyon, 69100, Villeurbanne, France; Université de Lyon, VetAgro Sup, 69280 Marcy l’Etoile, France; Unité Antibiorésistance et Virulence Bactériennes, Université de Lyon - ANSES Site de Lyon, Lyon, France

**Author notes:** Correspondence to Maria-Halima Laaberki. Contributed equally (Maria-Halima Laaberki, Xavier Charpentier).

## Abstract

With a great diversity in gene composition including multiple putative antibiotic-resistance genes, AbaR islands are potential contributors to multi-drug resistance in *Acinetobacter baumannii*. However, the effective contribution of AbaR to antibiotic resistance and bacterial physiology remains elusive. To address this, we sought to accurately remove AbaR islands and restore the integrity of their insertion site. To this end, we devised a versatile scarless genome editing strategy. We performed this genetic modification in two recent *A. baumannii* clinical strains: the strain AB5075 and the nosocomial strain AYE which carry AbaR11 and AbaR1 islands of 19.7 kbp and 86.2 kbp, respectively. Antibiotic susceptibilities were then compared between the parental strains and their AbaR-cured derivatives. As anticipated by the predicted function of the ORF of this island, the antibiotic resistance profiles were identical between the wild type and the AbaR11-cured AB5075 strains. In contrast, AbaR1 carries 25 ORFs with a predicted resistance to several classes of antibiotics and the AYE AbaR1-cured derivative showed restored susceptibility to multiple classes of antibiotics. Moreover, curing of AbaRs restored high levels of natural transformability. Indeed, most AbaR islands are inserted into the *comM* gene involved in natural transformation. Our data indicate that AbaR insertion effectively inactivates *comM* and that the restored *comM* is functional. Curing of AbaR consistently resulted in highly transformable, and therefore, easily genetically tractable strains. Emendation of AbaR provides insight into the functional consequences of AbaR acquisition.

## INTRODUCTION

*Acinetobacter baumannii* is responsible for healthcare-associated infections and concerns because of its intrinsic resistance to many antimicrobials and its ability to acquire resistance genes (1,2). A potential contributor to multi-drug resistance in *A. baumannii* is a genomic island, named AbaR, with a great diversity in gene content and carrying multiple putative antibiotic-resistance genes (3,4). The first report of AbaR described a large genomic island of 86 kbp in the epidemic *A. baumannii* strain AYE in 2006 (5). Since then, analysis of more than 3000 *A. baumannii* genomes deposited in the NCBI database revealed that AbaR are present in nearly 65% of them (4). With a transposon backbone, AbaR islands are considered as mobile genetic elements and present a modular structure (3,6–8). Although AbaR may contain up to 25 putative antibiotic-resistance genes, most AbaR carry three to four genes predicted to confer resistance mainly to aminoglycosides and tetracycline and with a median length of 20 kbp (4,9). Despite increasing genomic data, the actual contribution of AbaR-type islands to antibiotic resistance has not been fully evaluated. Deletion mutants of the AbaR23 island were previously obtained and suggested a paradoxical role of AbaR in increasing susceptibility to ciprofloxacin (9). However, deletions of AbaR23 was obtained by allelic replacement of AbaR23 with the *aacC1* cassette conferring resistance to gentamicin and confounded susceptibility testing against aminoglycosides. In addition, the original insertion site of AbaR was not restored which prevented analysis of the physiological consequence of the insertion of AbaR in the targeted gene. AbaR islands have been found to insert into 51 chromosome-borne loci, yet the overwhelming majority of AbaR islands are found inserted in the *comM* gene (4). This later gene encodes for a putative ATPase involved in natural transformation in a few transformable species including non-pathogenic *Acinetobacter baylyi* (10,11). Natural transformation allows bacteria to actively import exogenous DNA and, if the sequence of the imported DNA presents sufficient identity, it integrates the recipient cell’s chromosome by homologous recombination (12,13). Interestingly, the *comM* gene is not strictly essential for transformation but its mutation decreases the transformation efficiency by two orders of magnitude (10,11). *A. baumannii* and related species *A. nosocomialis* were described as transformable in 2013 (14,15). In *A. baumannii*, natural transformation appears as a conserved trait among clinical and non-clinical isolates of human and animal origin (14,16), providing the bacteria with a major route for acquisition of antibiotic resistance genes. In this study, we aim at assessing the contribution of AbaR islands to antibiotic resistance of two human clinical strains of the main Global Clone 1 bearing small and large AbaRs. We also analyzed the consequences of AbaR-mediated *comM* gene disruption on the transformation efficiency of these strains. We bring evidence that interruption of the *comM* gene by AbaR results in a decrease in natural transformation. Using genetic approaches, we further confirm a role for the *comM* gene in natural transformation in *A. baumannii*.

## MATERIALS AND METHODS

### Bacterial strains, typing and growth conditions

The bacterial strains used in this study are listed in Table S1. Unless specified, *A. baumannii* isolates and strains were grown in lysogeny broth (LB) (Lennox). All experiments were performed at 37°C. Antibiotic concentrations in selective media were 30 µg/mL for apramycin and 100 µg/mL for rifampicin.

### Construction of bacterial strains

All oligonucleotides used in this study for genetic modification are listed in supplementary Table S2. Gene disruptions were performed using overlap extension PCR to synthesize large chimeric DNA fragments carrying the selection marker flanked by 2 kbp-long fragments that are homologous to sequence flanking the insertion site. The PCRs were performed with a high-fidelity DNA polymerase (PrimeStarMax, Takara) (see section *Detailed protocol for chimeric PCR for mutation in A. baumannii* in the supplemental material). When used as template for the generation of PCR products, genomic DNA was extracted and purified using the Wizard® Genomic DNA purification kit (Promega) following the manufacturer’s guidelines. The AB5075 wild type strain was naturally transformed with an assembly PCR product to generate the AB5075 *comM::sacB_aac* strain. The AYE *comM::sacB_aac* was then obtained by transforming the AYE wild type strain with genomic DNA extracted from the AB5075 *comM::sacB_aac* strain. For both strains, transformants were selected on LB plates supplemented with apramycin and susceptibility to sucrose was verified on M63 plates supplemented with 10% sucrose. The AB5075 and AYE *comM::sacB_aac* strains were naturally transformed with an assembly PCR product (obtained from assembly PCR on wild type genomic DNA) to generate their -T and Δ*comM* derivatives. Transformants were selected on M63 plates supplemented with 10% sucrose and susceptibility to apramycin was verified on LB plates supplemented with apramycin. All chromosomal modifications were verified in transformants using colony PCR (cf. Table S2). Removal of AbaR and restoration of the *comM* gene were verified by Sanger sequencing.

The AB5075 rifampicin resistant (RifR) strain was generated by selecting spontaneous mutants after plating an overnight culture of the AB5075 wild type strain on plates supplemented with rifampicin.

### Antibiotic susceptibilities testing

Susceptibility of the strains to a panel of 15 antibiotics (ticarcillin, ticarcillin-clavulanic acid, piperacillin, piperacillin-tazobactam, ceftazidime, cefepime, meropenem, imipenem, gentamicin, tobramycin, amikacin, ciprofloxacin, tetracycline, aztreonam and sulfonamide) was evaluated by disc diffusion on Muller-Hinton agar (Biorad, France) following the CA-SFM 2013 recommendations (https://resapath.anses.fr/resapath_uploadfiles/files/Documents/2013_CASFM.pdf). Inhibition values were interpreted according to CA-SFM 2013 breakpoints for all antibiotics but aztreonam, for which *Pseudomonas* spp. breakpoints are given. To compare tetracycline susceptibility (as resistance marker of AbaR1) susceptibilities of wild type AYE and deleted AbaR1 mutant were compared by E-test (bioMérieux, France). The strain *Pseudomonas aeruginosa* CIP 7110 was used as control.

### Transformation assay

The method used in this study was described in (16) with the following modifications. After overnight incubation at 37°C on Lennox Broth agar, the strains were cultured for a few hours in 2 mL of Lennox Broth (LB) until an OD600nm of at least 1. The bacterial broths were then diluted to an OD600nm of 0.01 in PBS. Then, small aliquots of bacterial suspensions (5-10 µl) were mixed with DNA substrate. To measure transformation frequency, a final PCR concentration of 50 ng/µl was used. The mixtures (2.5 µl) were deposited on the surface of 1 ml of transformation medium (2% agarose D3 (Euromexdex), 5g/L of tryptone, 2,5g/L NaCl) /poured in 2-mL micro tubes and incubated overnight at 37°C. The next day, bacteria were recovered by resuspension in 300 µl of PBS. For genome edition experiments, the suspensions were spread on selective media (see *Construction of bacterial strains* section for details). To measure transformation frequency the bacterial suspensions were serial diluted and spread on LB agar plate without antibiotic or supplemented with rifampicin. All the transformation assays were performed on at least two separate experiments. On each occasion, at least three independent transformation reactions were conducted (three different transformation tubes). All the independent data points are plotted. As normality of the distribution of transformation frequency does not apply for transformation frequency analysis, non-parametric tests were performed (Mann-Whitney-Wilcoxon).

## RESULTS AND DISCUSSION

### Curing AbaR in A. baumannii using a scarless genome editing strategy

In order to cure *A. baumannii* of the AbaR island while restoring the original insertion site, we devised a scarless genome editing strategy. In contrast to other methods, the method does not require cloning, or prior genetic engineering of the target strain (17–19). Rather, it relies on the use of chimeric PCR products and takes advantage of natural transformation, a phenotypic trait exhibited by most isolates of *A. baumannii*. The method is based on a two-step selection (Fig. 1), first the introduction of a selectable/counter-selectable *sacB_aac* cassette at the loci of interest (Fig. 1B), then the replacement of the cassette by the desired sequence (Fig. 1C). First, a chimeric PCR product is assembled, consisting of the *sacB_aac* cassette bearing a selection marker (*aac(3)IV)* conferring resistance to apramycin, flanked by 2 kb-long sequences identical to the target region (Fig. 1B). This chimeric PCR product is introduced by natural transformation and *A. baumannii* transformants are selected using apramycin, an antibiotic to which most clinical isolates are sensitive. A second PCR product is assembled, still carrying the homologous flanking regions but whose sequence is designed to produce deletion, insertion or single-base mutation (deletion in Fig.1C). Genetic replacement and the loss of previously inserted *sacB* gene is then selected using sucrose resistance selection. We exemplified the method by deleting AbaR11 in strain AB5075. The strain AB5075 is an *A. baumannii* ST1 strain isolated from a bone infection in 2008 with multidrug resistance to eight antibiotic classes (aminoglycosides, fluoroquinolones, beta-lactams including penicillins, third and fourth generation cephalosporins, and carbapenems) (20). This strain bears a 19.7 kbp-long AbaR11 inserted in the *comM* gene with 19 predicted genes corresponding to five modules found in other AbaRs (Fig. 2A) (9,21,22). In a previous study, we identified the strain AB5075 as naturally transformable (16). Using natural transformation of PCR chimeric products, the AbaR island was replaced upon genetic recombination by the *sacB_aac* cassette (Fig. 2B). Then, the strain bearing the *sacB_aac* cassette in the *comM* gene was subjected to transformation with an assembly PCR encompassing the insertion region cured of AbaR thereby carrying an intact *comM* gene. Then we performed genetic analysis of sucrose-resistant and apramycin-susceptible clones to select an AbaR-cured strain (named AB5075-R) with a restored *comM* gene presumably encoding a full-length ComM protein (Fig. 2C).

**Figure 1.**
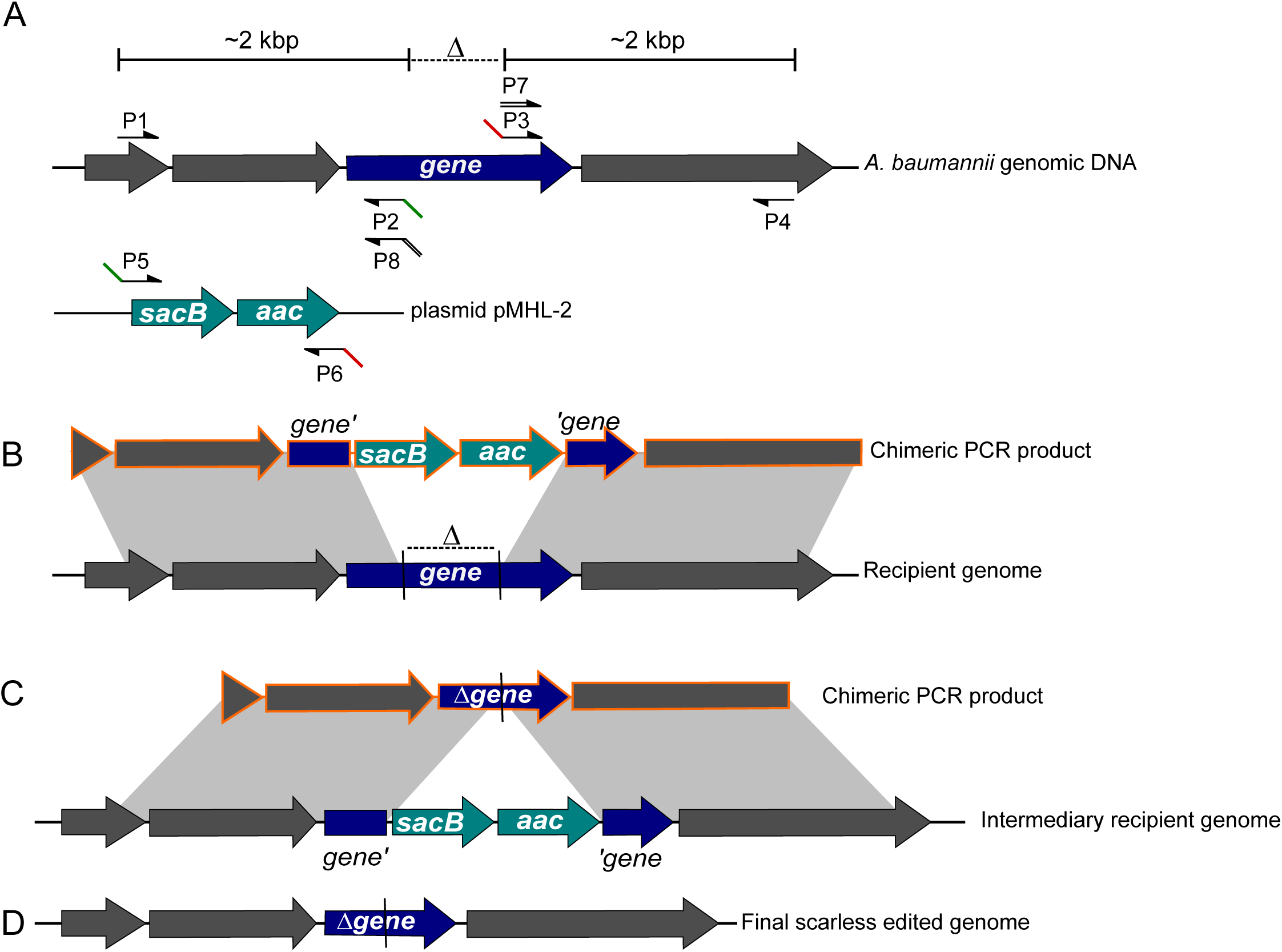
PCR-based genome editing of transformable *A. baumannii*. Schematic representations of steps for gene inactivation (dark blue gene) using natural transformation (A) Annealing sites for primers of PCR assemblies on their respective templates: either *A. baumannii* genomic DNA for amplifications of 2 kbp upstream and downstream of the deleted region (Δ) or the laboratory plasmid pMHL-2 that bears the counter-selectable cassette (light blue) with a *sacB* gene conferring sucrose sensitivity and the *aac(3)IV* gene encoding for resistance to apramycin. Note that primers P2, P3 and P7 share annealing sequences with primers P5, P6 and P8 respectively. (B) Chimeric PCR product (orange) obtained after two steps PCR assembly of products [P1 + P2] with [P5 + P6] and [P3 + P4]. This chimeric product bears the *sacB_aac* cassette flanked by two 2 kbp-long regions of homology with the recipient cell’s genome (grey areas). These homologous regions allow the recombination of the chimeric PCR product into the recipient cell’s genome resulting, upon apramycin selection of the transformants, in the intermediary genotype represented in C. (C) Chimeric PCR product obtained after two steps assembly PCR of products [P1 + P8] with [P7 + P4] bearing the deleted gene flanked by homologous regions (grey areas) allowing recombination and replacement of the counter-selectable cassette in the intermediary recipient genome. The transformants are selected based on their sucrose resistance resulting in the final scarless edited genome (D).

**Figure 2.**
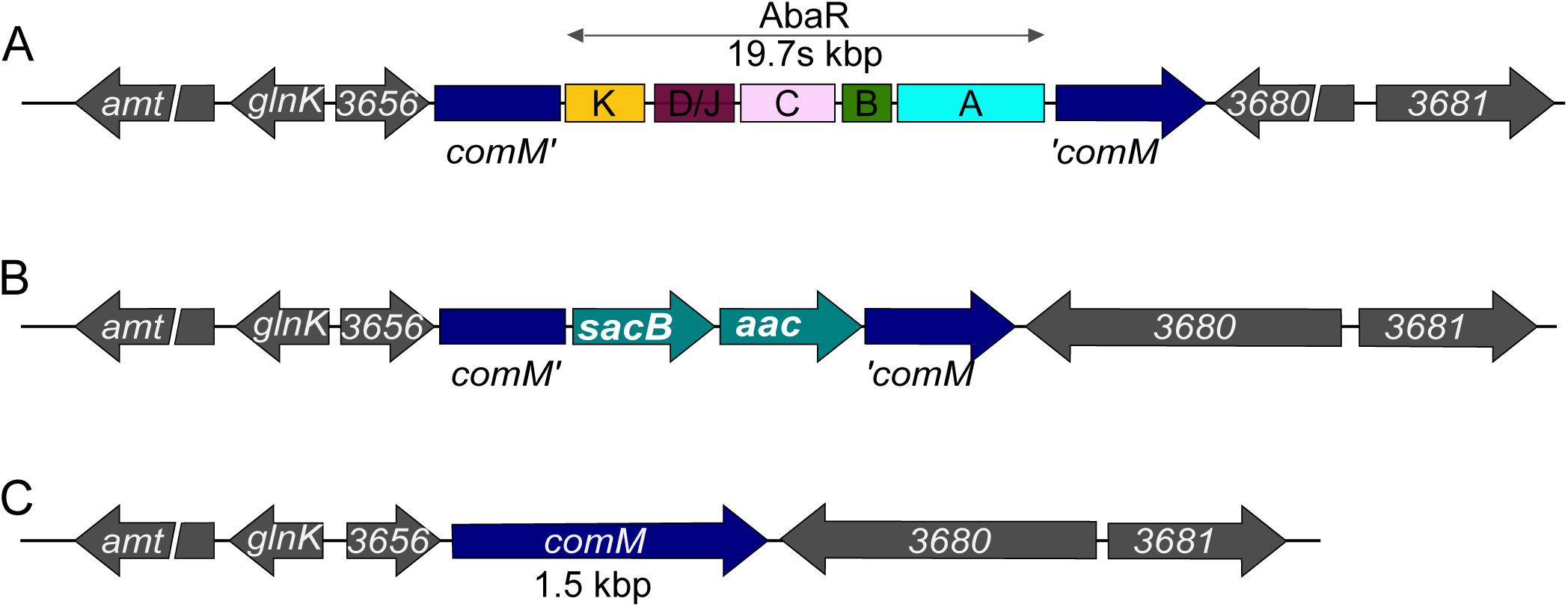
Genetic engineering of AbaR curing in *A. baumannii.* Schematic representations of (A) AbaR11 inserted in the *comM* gene in *A. baumannii* AB5075 genome. All the sequences and genes in grey color belong to the AB5075 WT chromosome, the interrupted *comM* gene is however highlighted in dark blue. For the AbaR island, modules are labeled following previous convention (9). For illustration purposes, scale of the AbaR island is not respected (B) Intermediary genetic construct with the counter-selectable cassette replacing the AbaR11 island. (C) Final genetic structure of the locus after restoring the *comM* gene integrity.

### Resistance profiles associated with AbaR curing in two clinically relevant MDR strains

We then sought to investigate the role of AbaR island in the multi-drug resistance profile of strain AB5075. With 19.7 kbp, AbaR11 falls in the median size of AbaRs (4). This AbaR contains 19 ORFs with, besides transposon-related proteins, seven ORFs predicted to confer resistance to metals, three of unknown function and one (*sup*) related to sulfonamide resistance encoding a putative sulfate permease present in the K-module found in most AbaRs (Fig. 2A). Antibiotic susceptibilities were thus compared between the AB5075 parental strain and its AbaR-cured derivative, denoted AB5075-T (Table 1). As anticipated, the resistance profiles to 15 antibiotics belonging to five antibiotic classes were identical between the wild type and its AbaR-cured derivative including to sulfonamides. We then investigated the contribution to antibiotic resistance of a larger AbaR found only in another *A. baumannii* ST1 strain, the clinical strain AYE resistant to aminoglycosides, ß-lactams (excluding imipenem, piperacillin-tazobactam, and ticarcillin-clavulanate), fluoroquinolones, and tetracycline and was of intermediate susceptibility to rifampicin (5,23). This strain carries the largest AbaR described so far, AbaR1, with 86 kbp-long, containing 25 genes putatively involved in resistance to antibiotic of several classes. By exploiting natural transformation of the strain AYE (16), we cured this strain of its AbaR1 island and compared the resistance profiles of the parental strain and its AbaR-cured AYE-T derivative (Table 1). In contrast to strain AB5075, curing the AbaR island from the strain AYE resulted restored susceptibility to aminoglycosides, tetracycline, sulfonamide and to some ß-lactams. A partial decreased resistance to ticarcillin (carboxypenicillin), ceftazidime (3GC), cefepime (4GC), aztreonam (monobactam) was also observed. The resistance *armamentarium* conferred by the 86 kbp-long AbaR1 may be attributed to the *aadB, aac6, aac3* and *aadDA1* genes for the resistance to amikacin, tobramycin and gentamicin (aminoglycosides), to the two *tetA-R* genes for tetracycline resistance and, to five copies of the *sulI* gene for the sulfonamide resistance (5). Partial loss in resistance to ticarcillin, ceftazidime, cefepime and aztreonam may be mainly attributed to loss of the *bla*_*VEB-1*_ gene and to a lesser extent the *bla*_*oxa-10*_ gene, this latter conferring resistance to all ß-lactams but carbapenems and extended spectrum cephalosporins (5,23). Only a partial loss of resistance to beta-lactams upon AbaR1 curing is likely explained by the additional resistance mechanisms of the AYE strain possess, in particular the over-expression of the *bla*ADC chromosomal cephalosporinase or from other genomic locations with the prediction of a new class A ß-lactamase on AYE chromosome (5,24). Contrarily to a previous report, we did not observe for both strains an increased resistance to fluoroquinolone of the AbaR-cured strains (9).

**Table 1.** Antimicrobial resistance profiles of *A. baumannii* strains AB5075 and AYE deleted of their AbaR island (-T) in comparison to the wild type strains (WT).

| Antibiotics<sup>a</sup> | Inhibition zone diameter in mm (category<sup>b</sup>) |  |  |  |
| --- | --- | --- | --- | --- |
|  | AB5075 WT | AB5075-T | AYE WT | AYE-T |
| AMK | 20 (S) | 20 (S) | 8 (R) | 21 (S) |
| GEN | 11 (R) | 12 (R) | 6 (R) | 19 (S) |
| TOB | 10 (R) | 10 (R) | 6 (R) | 18 (S) |
| CIP | 6 (R) | 6 (R) | 6 (R) | 6 (R) |
| PIP | 6 (R) | 6 (R) | 6 (R) | 6 (R) |
| TZP | 6 (R) | 6 (R) | 14 (R) | 15 (R) |
| TIC | 6 (R) | 6 (R) | 6 (R) | 19 (I) |
| TIM | 6 (R) | 6 (R) | 15 (I) | 18 (I) |
| CAZ | 6 (R) | 6 (R) | 6 (R) | 11 (R) |
| FEP | 6 (R) | 6 (R) | 6 (R) | 16 (I) |
| IMP | 6 (R) | 6 (R) | 28 (S) | 28 (S) |
| MEM | ND | ND | 20 (S) | 22 (S) |
| ATM | 6 (R) | 6 (R) | 6 (R) | 11(R) |
| SUF | 6 (R) | 6 (R) | 6 | 23 |
| TET | 18 (S) | 18 (S) | ND | ND |
| TET (E-test)<sup>c</sup> | ND | ND | 128 | 8 |

<sup>a</sup> Abbreviations. AMK: Amikacin, GEN: Gentamicin, TOB: Tobramycin, CIP: Ciprofloxacin, PIP: Piperacillin, TZP: Piperacillin + tazobactam, TIC: Ticarcillin, TIM: TIC + clavulanic acid, CAZ: Ceftazidime, FEP: Cefepime, IMP: Imipenem, MEM: Meropenem, ATM: Aztreonam, TET: Tetracycline, SUF: sulfonamide.
<sup>b</sup> Categorization (R: resistant, S: susceptible) is detailed in the materials and methods section.
<sup>c</sup> E-test strips were used to determine the MICs of tetracycline for the strains AYE and AYE-T.
ND: Note Done.

### Curing of AbaR and repair of the comM gene result in increased transformation ability

Among the 51 chromosome-borne insertion loci, AbaR islands are predominantly inserted in the *comM* gene (4). In the non-pathogenic species *A. baylyi*, a mutation of this latter gene is associated with a reduced transformability (10). We therefore pursued by investigating the role of *comM* in *A. baumannii* and the physiological consequence of AbaR-island disruption of this gene. To do so, we performed transformation assays comparing the levels of transformation efficiency between the wild type and AbaR-cured strains (Fig. 3). For both clinical strains, curing AbaR and thus restoring the *comM* gene increased significantly the ability to acquire a genetic marker with a mean increase of about a 1000-fold in strain AB5075-T (Fig. 3A) and of about 300-fold in strain AYE-T (Fig. 3B). With one bacterium out of 100 that performed natural transformation, strain AB5075-T presented the highest transformation ability (Fig. 3A). Using the same genome-editing strategy as depicted in Fig. 1, we generated markerless deletions of the *comM* gene (Δ*comM*) in both *A. baumannii* clinical strains. Consistently, deletion of the *comM* gene in both strains resulted in transformation levels comparable to levels observed for the wild type strains bearing a genuine insertion in the *comM* gene by the AbaR island (Fig. 3A and B). Thus, prior to acquisition of AbaR, both strains were transformable at higher levels, and the subsequent disruption generates a transformation defects that is consistent with previous observations made in other species or genus (10,11). The data show that the AbaR-inactivated *comM* gene, once cured of AbaR, is still functional. This suggests that it has not been affected by mutational drifts, confirming that acquisition of AbaR occurred relatively recently (22).

**Figure 3.**
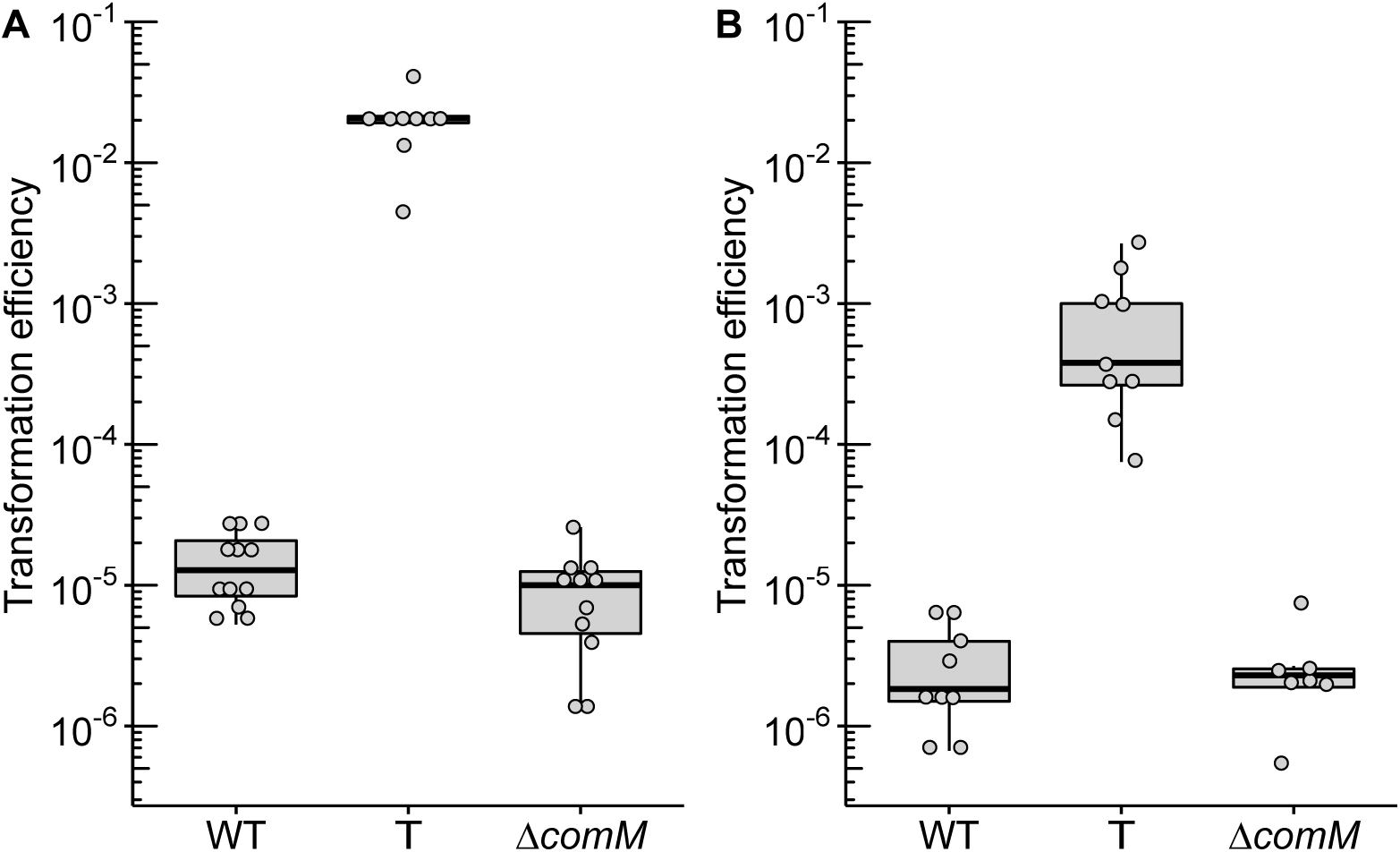
Transformation efficiency of *A. baumannii* strains AB5075 (A) and AYE (B) of a 3,8 kbp-long PCR product carrying a mutation in the *rpoB* gene conferring resistance to rifampicin. For both strains, either the wild type, their “T” (ΔAbaR / repaired *comM*) or Δ*comM* derivatives were tested. At least seven independent transformation assays were performed on at least two separate occasions represented by separate dots and corresponding boxplots. All results were above the limit of detection (10^−8^). Pairwise comparisons using the non-parametric Mann-Whitney-Wilcoxon test (two tailed) gave at least *p*-values <0.005 between transformation efficiencies obtained for the wild type/Δ*comM* strains and the T-derivatives.

The fact that acquisition of the AbaR element in the *comM* gene alters the natural transformability of *A. baumannii* and its capacity for further acquisitions by natural transformation is intriguing. The significance of lower transformability is not known but theoretical work suggest that intermediate transformation rates allows bacteria to maximize fitness in fluctuating, stressful environments such as repeated antibiotic stress (25). Our results then suggest that acquisition of AbaRs, and associated resistance, provide a short-term benefit in environment with constant exposure to antibiotics but may ultimately lead to an evolutionary dead end.

## CONCLUSION

By exploiting natural transformation, we setup a genetic system to easily edit the genome of *A. baumannii*. We used this strategy to perform small gene deletion but also to cure large genomic islands in two MDR clinical *A. baumannii* strains. Moving beyond predicted resistance, this system allowed experimental determination of the antibiotic resistance conferred by the islands. We conclude that acquisition of AbaR islands may increase *A. baumannii* strains resistance to antibiotics but it comes at the cost of decreasing the cell ability to perform natural transformation by disrupting the *comM* gene. Thus, acquisition of AbaR may impact genome evolution of *A. baumannii*. From an applied perspective, the simultaneous deletion of AbaR and restoration of the *comM* function in two clinically relevant strains made them even more genetically amenable, blazing the trail for mutational analysis of these model *A. baumannii* strains.

## Acknowledgments

We thank the LABEX ECOFECT for building an “ecofectian” community.

