## Supplemental material for "Scarless removal of large resistance island AbaR results in antibiotic susceptibility and increased natural transformability in *Acinetobacter baumannii*"

**Table S1: *Acinetobacter baumannii* strains used in this study**

**Table S2: primers used in this study**

**Detailed protocol of chimeric PCR for mutation in *A. baumannii***

SUPPLEMENTARY MATERIALS

**Table S1: *Acinetobacter baumannii* strains used in this study**

| Strain name | Genotype | Description/Phenotype | Reference |
|---|---|---|---|
| AB5075 WT | *A. baumannii* AB5075 wild type | | (1) |
| AB5075 *comM*::*sacB_aac* | *A. baumannii* AB5075 *comM*::*sacB_aac* | replacement of the AbaR island with the *sacB_aac(3)* cassette, resistant to apramycin and sensitive to sucrose | this study |
| AB5075-T | *A. baumannii* AB5075 $\Delta$AbaR11 | deletion of the AbaR11 island and a repaired *comM* gene, highly transformable strain | this study |
| AB5075 $\Delta$*comM* | *A. baumannii* AB5075 $\Delta$*comM* | markerless 231 bp deletion of the *comM* gene | this study |
| AB5075 Rif$^{R}$ | *A. baumannii* AB5075 Rif$^{R}$ | spontaneous *rpoB* mutant; rifampicin resistant | this study |
| AYE WT | *A. baumannii* AYE wild type | wild type | (2) |
| AYE *comM::sacB_aac* | *A. baumannii* AYE *comM::sacB_aac* | strain with a replacement of the AbaR island with the *sacB_aac(3)* cassette, resistant to apramycin and sensitive to sucrose | this study |
| AYE-T | *A. baumannii* AYE $\Delta$AbaR1 | strain with a deletion of the AbaR1 island and a repaired *comM* gene, highly transformable and antibiotic susceptible strain | this study |
| AYE $\Delta$*comM* | *A. baumannii* AYE $\Delta$*comM* | markerless 231 bp deletion of the *comM* gene | this study |


1. Hujer KM, Hujer AM, Hulten EA, Bajaksouzian S, Adams JM, Donskey CJ, et al. Analysis of Antibiotic Resistance Genes in Multidrug-Resistant Acinetobacter sp. Isolates from Military and Civilian Patients Treated at the Walter Reed Army Medical Center. Antimicrob Agents Chemother. 2006 Dec;50(12):4114–23.
2. Fournier P-E, Vallenet D, Barbe V, Audic S, Ogata H, Poirel L, et al. Comparative genomics of multidrug resistance in Acinetobacter baumannii. PLoS Genet. 2006 Jan;2(1):e7.

**Table S2: primers used in this study**

| Genetic construct | Name* | Sequence 5' to 3' | Template; Annealing site |
|---|---|---|---|
| Chromosomal modifications using overlap extension PCR | | | |
| *comM::sacB_aac* | mlo-2 (P1) | GCCTGGAATTGTCATCAACAGAACC | AB5075 and AYE genomic DNA; *amt* gene = 2kb upstream of the *comM* gene |
| | mlo-3 (P2) | **GGCCCAATTCGCCCTATAGTGAGTCG**ATTGCTGATGCTGTATGGTGGGG | AB5075 and AYE genomic DNA; 5' *comM* gene and adapter sequence reverse-complement to primer mazF-K7-F (bold and orange) |
| | mlo-4 (P3) | **GGGTTTGCTCGGGTCGGTGGCATATG**CAATGAATCCTTGTCCATGTGGG | AB5075 and AYE genomic DNA; 3' *comM* gene and adapter sequence reverse-complement to primer mlo-1 (bold and blue) |
| | mlo-5 (P4) | GCATGTCTTGCTGTGCGTGTTGC | AB5075 and AYE genomic DNA; *ABUW_3681* gene = 2kb downstream of the *comM* gene |
| | mazF-K7-F (P5) | **CGACTCACTATAGGGCGAATTGGGCC**GCTTTCCAGTCGGGAAACCTG | pMHL-2 (Godeux et al., 2018) ; upstream *sacB* gene and adapter sequence (bold) |
| | mlo-1 (P6) | **CATATGCCACCGACCCGAGCAAACCC**CGCCAGGGTTTTCCCAGTCACGAC | pMHL-2 (Godeux et al., 2018) ; dowstream *aac(3)* gene and adapter sequence (bold) |
| ΔAbaR | mlo-6 (P7) | ACCGCCCCCAACCAGTGCAATTG | AB5075 and AYE genomic DNA ; *comM* at AbaR insertion locus (annealing with mlo-7 is underlined) |
| | mlo-7 (P8) | CAATTGCACTGGTTGGGGGCGGTTCACACCCCAAACCGGGTGAAATTAC | AB5075 and AYE genomic DNA ; *comM* at AbaR insertion locus (annealing with mlo-6 is underlined) |
| Δ*comM* | asg-75 (P7') | AATTTCTAGTGCACGCCG | AB5075 and AYE genomic DNA; 5' *comM* gene (annealing with asg-76 is underlined) |
| | asg-76 (P8') | CGGCGTGCACTAGAAATTCCAAACCGGGTGAAATTAC | AB5075 and AYE genomic DNA; 3' *comM* gene (annealing with asg-75 is underlined) |
| Primers for control using colony PCR and for sequencing of the *comM* locus | | | |

| | | | |
|---|---|---|---|
| | mlo-80 | CGGATCTTCGATGCTGGC | AbaR1 from AYE strain |
| | mlo-84 | GCAACGATGTTACGCAGC | AbaR1 from AYE strain |
| | comM-For | CCACAATGGAACAAGAAGATGTCT | 5’ of the *comM* gene |
| | comM-Rev | TTAAGAGTGATTACCTCGATAAGA | 3’ of the *comM* gene |
| | mlo-29 | TCATGAGCTCAGCCAATCGACTGG | End of ApraR cassette |
| | asg-61 | CCACAGAATGATGTCACG | 5’ of the *aac* gene |
| PCR product carrying the *rpoB* mutation as substrate for natural transformation | | | |
| | mlo-104 | AAGATATCGGTCTCCAAGC | AB5075 genomic DNA; *rpoB* gene |
| | mlo-105 | AGTACGGCCTTCAACGTCAT | AB5075 genomic DNA; *rpoB* gene |

*as referred in Fig.1

**Detailed protocol of chimeric PCR for mutation in *A. baumannii***

Two rounds of PCR are required to produce chimeric PCR products: a first PCR to add overlapping extensions to the DNA fragments and a second to assemble the first DNA fragments.

1. **First PCR (Overlapping extension PCR)**

*PCR mix:*

| | |
|---|---|
| PCR mix | 25µL reaction |
| DNA template [20ng to 100ng/µL] | 1µL |
| Primer F 10µM | 0.5µL |
| Primer R 10µM | 0.5µL |
| PrimeSTAR premix (Takara) | 12.5µL |
| H2O qsp 25µL | 10.5µL |

*PCR cycling conditions :*

Initial denaturation 98°C- 2 min.

Denaturation 98°C - 20 sec.
Annealing **55**°C – 20 sec.
Extension 72°C – 40 min.
30 cycles

Final extension 72°C – 5 min.

Then proceed to PCR purification after migration of the whole PCR reactions in buffered agarose gel stained with SYBR Safe (Thermofisher). The bands at the correct size are excised from the gel above a blue-light transilluminator (this is critical) and DNA extracted following instructions of the manufacturer (Omega Bio-Tek/VWR).

Alternatively, the PCR products are purified using magnetic beads following manufacturer's protocol (Ampure XP /Beckman Coulter) after a direct DpnI digestion of the genomic DNA in the PCR mix.

The purified PCR products are then analyzed (5µL) on agarose gel stained with ethidium bromide. Concentration should be at least 30ng/µL for each PCR products and a single band obtained.

**2. Second PCR (assembly PCR)**

This second round of PCR assembles the three PCR products with a ratio of 1/2/1 for PCR[P1 + P2] / PCR[P5 + P6] / PCR[P3 + P4] in the PCR mix using the P1 (forward primer) and P4 (reverse primer).

*PCR mix:[DNA concentration for example]*

| PCR Assembly | 50µL reaction |
|---|---|
| PCR[P1 + P2] [100ng/µL] | 1 |
| PCR[P5 + P6] [30ng/µL] | 6 |
| PCR[P3 + P4] [100ng/µL] | 1 |
| Primer P1 10µM | 1 |
| Primer P4 10µM | 1 |
| PrimeStar premix (Takara) | 25 |
| H2O | 15 |

Note that the total amount of PCR products is close to 500ng for a 50µL reaction, this can be up to 800ng-1µg, without exceeding a total of 10µL of first PCR products for a 50µL PCR reaction.

*PCR cycling conditions :*

Initial denaturation 98°C- 2’

Denaturation 98°C -20”<br>Annealing 55°C – 20”<br>Extension 72°C – 1’ 30”<br>30 cycles

Final extension 72°C – 5’

Then proceed to PCR purification either after migration in buffered agarose gel stained using SYBR Safe (Thermofisher) followed by gel extraction (Omega Bio-Tek/VWR) or alternatively using Ampure XP beads (Beckman Coulter).
